# Cellular phenotyping of hippocampal progenitors exposed to patient serum predicts conversion to Alzheimer’s Disease

**DOI:** 10.1101/175604

**Authors:** Aleksandra Maruszak, Tytus Murphy, Benjamine Liu, Chiara de Lucia, Abdel Douiri, Alejo J Nevado, Charlotte E Teunissen, Pieter Jelle Visser, Jack Price, Simon Lovestone, Sandrine Thuret

## Abstract

The generation of new neurons persists into adulthood in the human hippocampus and can be modulated by the circulatory systemic environment. Hippocampal neurogenesis is important for learning and memory and is altered in Alzheimer’s Disease (AD). Evaluating the hippocampal neurogenic process during disease progression could therefore identify neurogenesis as an important target for AD prevention and intervention as well as a biomarker for early disease detection. In this study, we used a human hippocampal progenitor cell line to design an in vitro assay evaluating over time the neurogenic impact of the systemic milieu (i.e. serum) of individuals with mild cognitive impairment (MCI) as they either converted to AD or remained cognitively stable. Cells were exposed to serum collected over several years from the same patients. Cellular phenotyping and linear mixed effects models for repeated measures revealed that decreased proliferation, increased apoptotic hippocampal progenitor cell death and increased hippocampal neurogenesis characterized progression from MCI to AD. Using stepwise logistic regression and machine learning we show that these cellular readouts for the baseline serum sample and years of education of the patient are significant predictors of conversion from MCI to AD, already 3.5 years before AD clinical diagnosis. Finally, serum proteomic analyses indicated pathways linked to the cellular readouts distinguishing MCI to AD converters from non-converters. The proposed assay is thus not only promising for AD pre-clinical diagnosis, but it also provides a proxy into temporal changes of the hippocampal neurogenic process during disease progression.

**One Sentence Summary:** In this study, we demonstrate for the first time that the systemic environment (i.e. blood serum) of mild cognitively impaired patients differentially alters human hippocampal progenitor cell fate to predict conversion to Alzheimer’s Disease up to 3.5 years before clinical diagnosis.

## Introduction

Alzheimer’s disease (AD) is a progressive neurodegenerative condition without any effective treatment options. While researchers have traditionally focused on examining methods to avert neuronal loss in AD, adult new-born neuron generation, i.e. neurogenesis, is emerging as a target for prevention and therapeutic interventions as well as a potential biomarker for early disease detection. Human neurogenesis is a lifelong process, persisting in adulthood predominantly in the subgranular zone (SGZ) of the hippocampal dentate gyrus (*1*). The SGZ creates a permissive microenvironment, the so-called neurogenesis niche, for hippocampal progenitor cells to proliferate and differentiate (*2*). The niche provides cues for the survival and fate of the hippocampal progenitor cells, determining whether they will divide and generate neuroblasts. The neuroblasts migrate to the granule cell layer and differentiate into dentate granule neurons, which will project dendrites into the molecular layer of the hippocampus and axons toward the CA3 region of the hippocampus (*1, 3*).

The process of hippocampal neurogenesis (HN) is fundamental for hippocampal-dependent learning and memory (*4*). Therefore, HN dysfunction appears especially relevant to dementias and in particular to AD. The hippocampus is indeed one of the first affected brain structures in AD and its atrophy is associated with memory and learning impairment (*5, 6*). There is extensive evidence showing that altered HN is one of the early AD hallmarks (*7, 8*), although neither rodent models of AD studies nor human autopsy analyses have been in agreement and therefore collectively failed to provide conclusive information with regard to directionality and magnitude of these changes (*7, 8*). Moreover, there have been no studies looking at longitudinal changes in HN during AD progression. This approach is partially hampered by the fact that so far there are no neuroimaging techniques to visualise HN in the living human brain. However, an insight into the temporal sequence of HN changes over AD progression would enable the design and evaluation of interventions aimed at slowing down the disease progression or maintaining cognitive functions in the elderly.

The neurogenic niche comprises not only hippocampal progenitor cells but also their progeny -neurons, glia, endothelial cells and extracellular matrix integrating cell-to-cell contacts and cell signaling, all localized around blood vessels (*9*). Because the niche is highly vascularized, it allows potential communication with the systemic environment, a feature critical in the context of aging (*10-12*). Parabiosis experiments, in which the circulatory systems of two mice are surgically conjoined, demonstrated that blood from young mice has a cognitive rejuvenating effect on the old animals, which benefited from improved HN (*10, 13, 14*). Injecting plasma from the young to the old mice had similar effect and increased HN (*10, 13*). These studies highlighted for the first time the direct ability of the systemic environment in modulating HN. Moreover, several systemic interventions (i.e. drugs, exercise, diet) have been shown to modulate HN (*15-17*) and some, such as exercise (*18*) and particular diet components to decrease AD risk (*7, 19*).

The considerable abnormalities in the blood-brain barrier (BBB) in the hippocampi of AD patients (*20*) create a gateway for an altered exchange of nutrients, inflammatory molecules, drugs, hormones, growth factors, and other blood-derived factors, affecting the brain milieu and resulting in altered HN. Using dynamic contrast-enhanced MRI, Montagne et al. demonstrated that BBB damage in aging starts in CA1 and DG, and these changes are significantly more pronounced in mild cognitive impairment (MCI), the early symptomatic stage of AD (*20*). These findings suggest a route whereby the systematic environment might alter neurogenesis.

MCI is a heterogeneous condition and does not constitute a precise diagnosis as not all individuals with MCI develop AD, or indeed any neurodegenerative disease. Annual conversion rate to AD of individuals diagnosed with MCI (10-15% in clinical studies; 5-10% in population studies) is higher than of elderly healthy persons (1-2%) (*21-24*). Therefore, an accurate estimation of individual’s likelihood of conversion could be used to target interventions to treat or prevent disease progression, and applied in the design of clinical trials.

The consensus is that any putative AD-modifying therapies would be most beneficial if administered before the clinical onset of dementia or during the preclinical stage. Without an accurate predictor of the risk of conversion of MCI to AD, therefore, pharmacological and lifestyle interventions cannot be correctly targeted.

While the majority of AD research has utilized cross-sectional data, evidence from the Dominantly Inherited Alzheimer Network (DIAN) study indicates that longitudinal analyses provide a more accurate picture of disease progression (*25, 26*). Owing to the longitudinal, within-person assessments performed in the framework of the DIAN study, a temporal separation between first amyloid-β accumulation and later hippocampal atrophy could be established (*26*). This suggested that between these two pathological events there is room for neuroprotective mechanisms to preserve brain function. We believe that changes in HN level might be acting as compensatory response to the early pathological AD events. By utilising human hippocampal progenitor cells and longitudinal serum samples from MCI patients who either converted to AD (MCI converters) or remained cognitively stable (MCI non-converters), we aimed to establish the role of the human systemic environment during disease progression in an in vitro model of human HN (Fig.1). We also sought to determine if our HN assay could be used in predicting conversion from MCI to AD. Our study presents the first attempt to look at human hippocampal neurogenesis as a dynamic process and to apply it to prediction of conversion to AD.

**Fig. 1.**
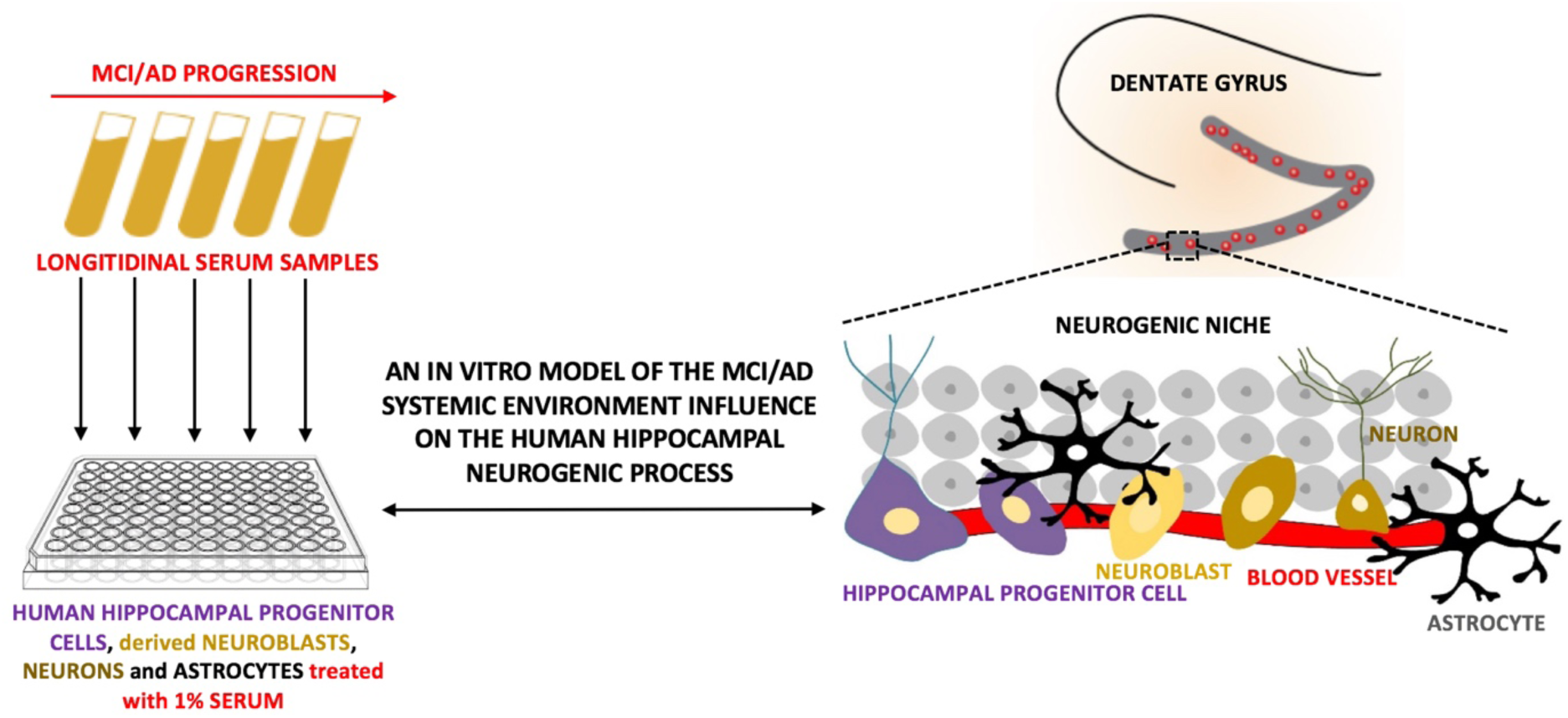
The proposed in vitro model to study the role of the systemic environment on the human hippocampal neurogenenic process in the context of Mild Cognitive Impairment (MCI) and Alzheimer’s disease (AD). Hippocampal neurogenesis is regulated by a complex microenvironment of hippocampal progenitor cells, i.e. neurogenic niche, composed not only of hippocampal progenitor cells but also neurons, glia and blood vessels. Recently, parabiosis studies demonstrated the fundamental role of the blood-derived factors (systemic environment), delivered to the niche by its rich vasculature, in modulating hippocampal neurogenesis. Our assay, in which we subject a human hippocampal progenitor cell line at different stages of proliferation and differentiation to 1% longitudinal patients’ serum during disease progression, aims to model in vitro the role of the systemic environment on the hippocampal neurogenic process, during AD progression.

## Results

### Effect of longitudinal serum samples from MCI converters on proliferation, differentiation and cell death of human hippocampal progenitor cells

To investigate the role of the human systemic environment (i.e. serum) on HN during AD progression, the HPC0A07/03C human hippocampal progenitor cell line (*15, 27*) was exposed to 1% serum collected up to 6 times for up to 6 years from the same patient progressing or not to AD. Fifty-six patients initially diagnosed with MCI were individually followed up. Thirty-eight of 56 MCI patients later developed dementia due to AD (MCI converters), whereas 18 did not progress either to AD or other disease (MCI non-converters) (Fig.2A). Serum was added to the cell culture during both proliferation and differentiation (Fig.2B). Cellular phenotyping with all sequential serum samples was characterized by immunohistochemistry and high content cellular imaging for proliferation, cell death and neurogenesis.

**Fig. 2.**
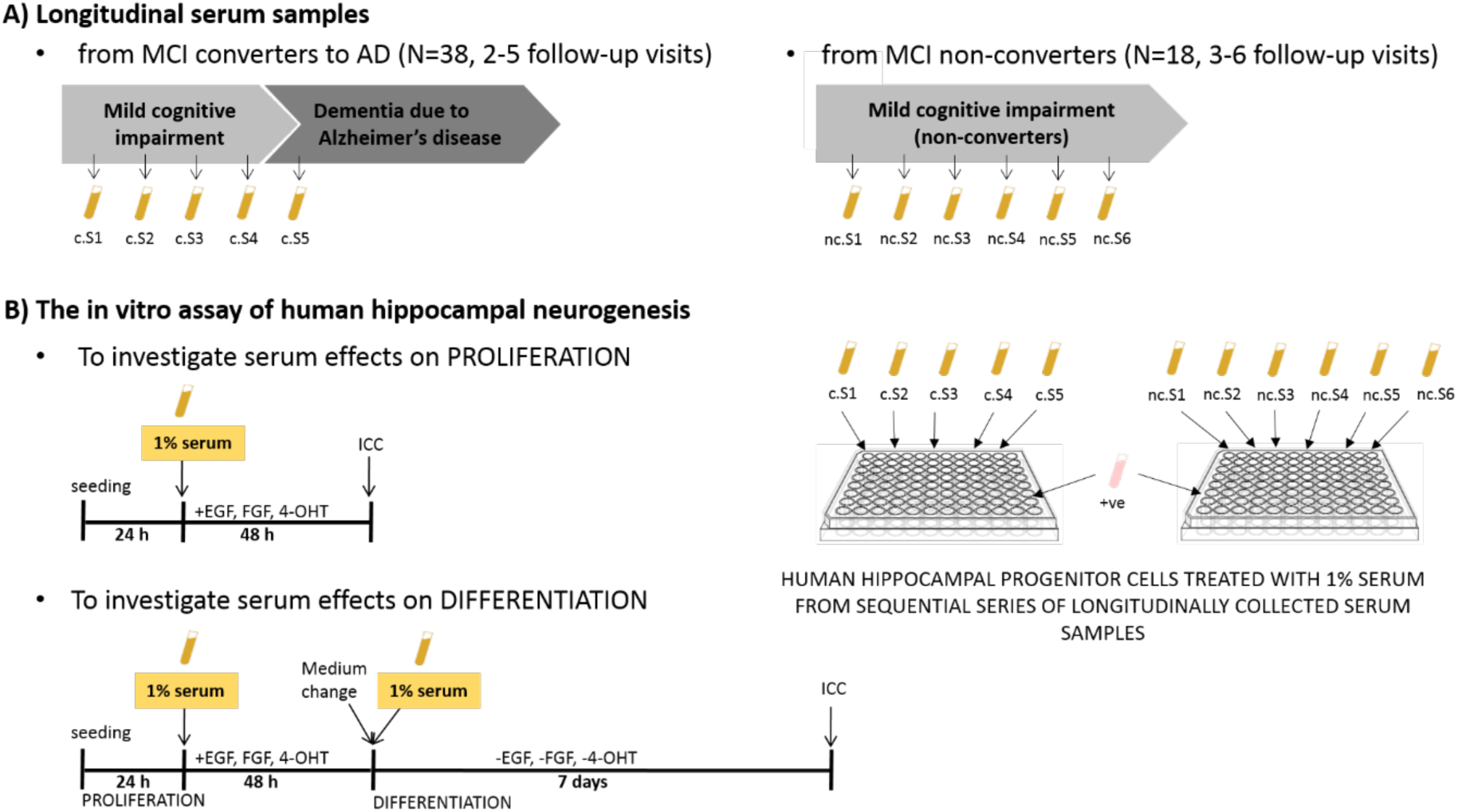
Outline of the patients’ sample collection (A) and the cellular assay (B). **A)** Longitudinal serum samples were collected during annual follow-up visits from 56 study participants diagnosed with MCI at baseline. 38 converted to AD, 18 remained cognitively stable. A minimum of three serum (S) samples, collected at annual assessment, was required for MCI non-converters (indicated with letters “nc.S” and a number corresponding to the visit), and two samples, one prior to conversion and one post-conversion, for MCI converters (indicated with letters “c.S” and a number corresponding to the visit). **B)** The in vitro assay of the human hippocampal neurogenic process to model the systemic environment influence on human hippocampal neurogenesis in the context of AD -overview of the proliferation and differentiation conditions of the HPC0A07/03C cell line in medium supplemented with 1% serum collected during sequential follow-up visits of MCI converters and non-converters. Proliferation medium comprised 4-hydroxytamoxifen (4-OHT), epidermal growth factor (EGF) and basic fibroblast growth factor (FGF). 24 hours after seeding, cell medium was replaced and supplemented with 1% serum. HPC0A07/03C were left to proliferate for further 48 hours. To analyse serum effects on proliferation, after 72 hours in culture, the cells were fixed in 4% PFA and next subjected to immunocytochemistry (ICC) labelling. In order to analyse the effects of serum on differentiation, 1% serum had to be present in cell medium during both proliferation and differentiation. Serum was added only once during each stage. After 72 hours of proliferation, medium was removed and replaced by one devoid of EGF, FGF, 4-OHT and supplemented with 1% serum. After a 7 day-differentiation the HPC0A07/03C cells were fixed and subjected to ICC. Medium supplemented with 1% serum was never changed during either proliferation or differentiation stage. Control conditions consisted of medium + ve (1:100 PenStrep, Thermo Fisher Scientific). Proliferation was evaluated with Ki67, apoptotic cell death with Cleaved Caspase 3 (CC3), neuroblasts with doublecortin (Dcx) and neurons with microtubule-associated protein 2 (Map2) labelling. Each experiment was run in three biological replicates (cells of three different passage numbers) and for each biological replicate there were technical triplicates.

First, using the longitudinal serum samples from the MCI converters and the presented assay, we modelled the effect of systemic environment on HN from baseline diagnosis until conversion to AD. We sought to investigate if disease progression could be monitored using a serum sample from each follow-up visit and our in vitro assay. We obtained a neurogenesis readout corresponding to each follow-up visit and performed modelling of the intra-and inter-individual differences in the systemic effects on human HN. Mixed-effects models for repeated measures were fitted to identify factors affecting trajectories of HN changes. Since age did not significantly predict HN level (p>0.05), we used another measure enabling longitudinal analysis -time to conversion, i.e. time in years to the clinical diagnosis of dementia due to AD. Because the last serum sample used in our study was obtained at the time of conversion to AD, it was assigned the value of 0, and time before conversion assigned negative values (i.e. one year before conversion is −1 year).

### Decreased proliferation and increased apoptotic hippocampal progenitor cell death characterize progression from MCI to AD

The model with random intercept predicts that over the disease progression there is an increase in the number of hippocampal progenitor cells (p=0.002, Fig.3A and B, Table 1). This is not related to an increase in proliferation, as conversely, decreasing level of proliferation (%Ki67+ cells, p<0.0001, Fig.3A and D, Table 1) of hippocampal progenitor cells across all analysed time points was detected. Time to conversion was the only significant predictor of proliferation and cell number in both random intercept models (see Table 1) for average cell count and %Ki67+ cells.

**Fig.3.**
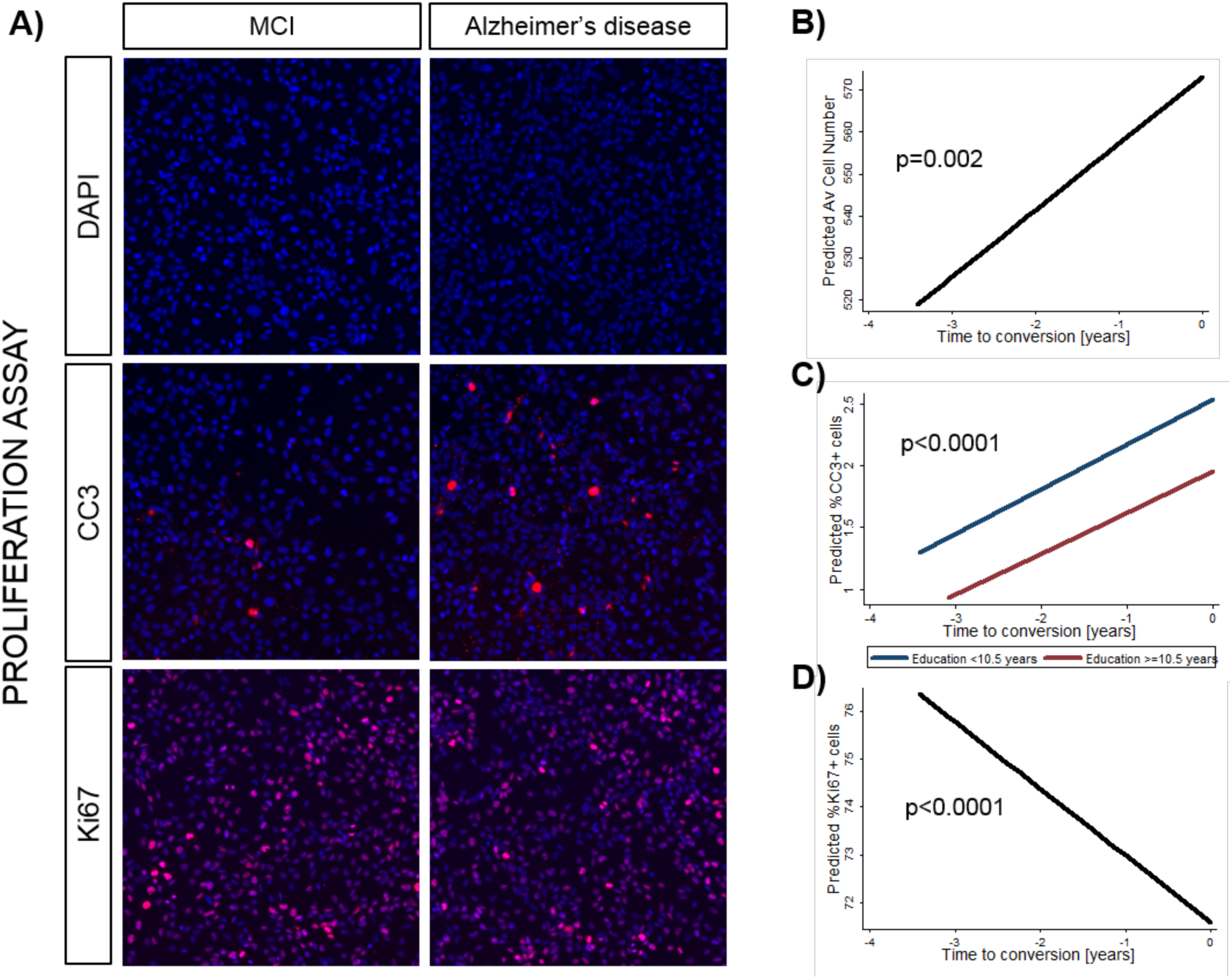
Exposure to 1% serum from MCI converters leads to decreased proliferation and increased cell death of the HPC0A07/03C cells. **(A)** Representative images of human hippocampal neurogenesis when the HPC0A07/03C cells are treated with serum from the same individual –on the left with serum sample from one year before conversion (MCI) and on the right with serum sample taken at the conversion to AD. Increase in cell number (DAPI), decrease in proliferation (Ki67 labelling) and increased apoptotic cell death (Cleaved Caspase 3, CC3 labelling) of hippocampal progenitor cells were detected. **(B-D)** The graphs present results of fitting mixed-effects models for repeated measures for MCI converters. For the average cell number and Ki67 models with random intercept were the best fit. For CC3 random slope. Trajectories of hippocampal neurogenesis as a function of time to conversion are shown. On the x-axis conversion event is depicted as year=0 and time before conversion is assigned negative values. Longitudinal serum samples from MCI converters increased cell count **(B)**, cell death (%CC3+ cells) **(C)** and decreased proliferation (%Ki67+ cells) **(D)** of the HPC0A07/03C cells. All fitted regression lines were calculated from models presented in Table 1.

**Table 1.**
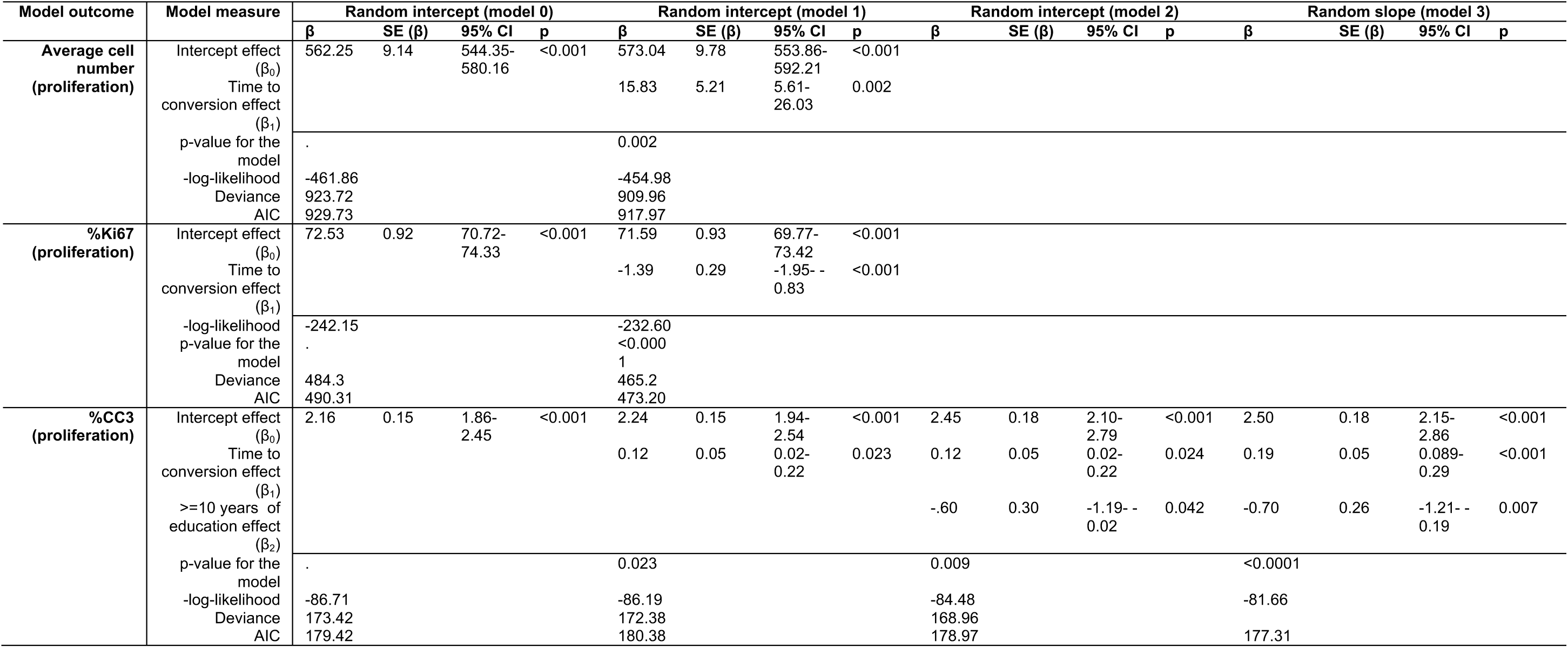
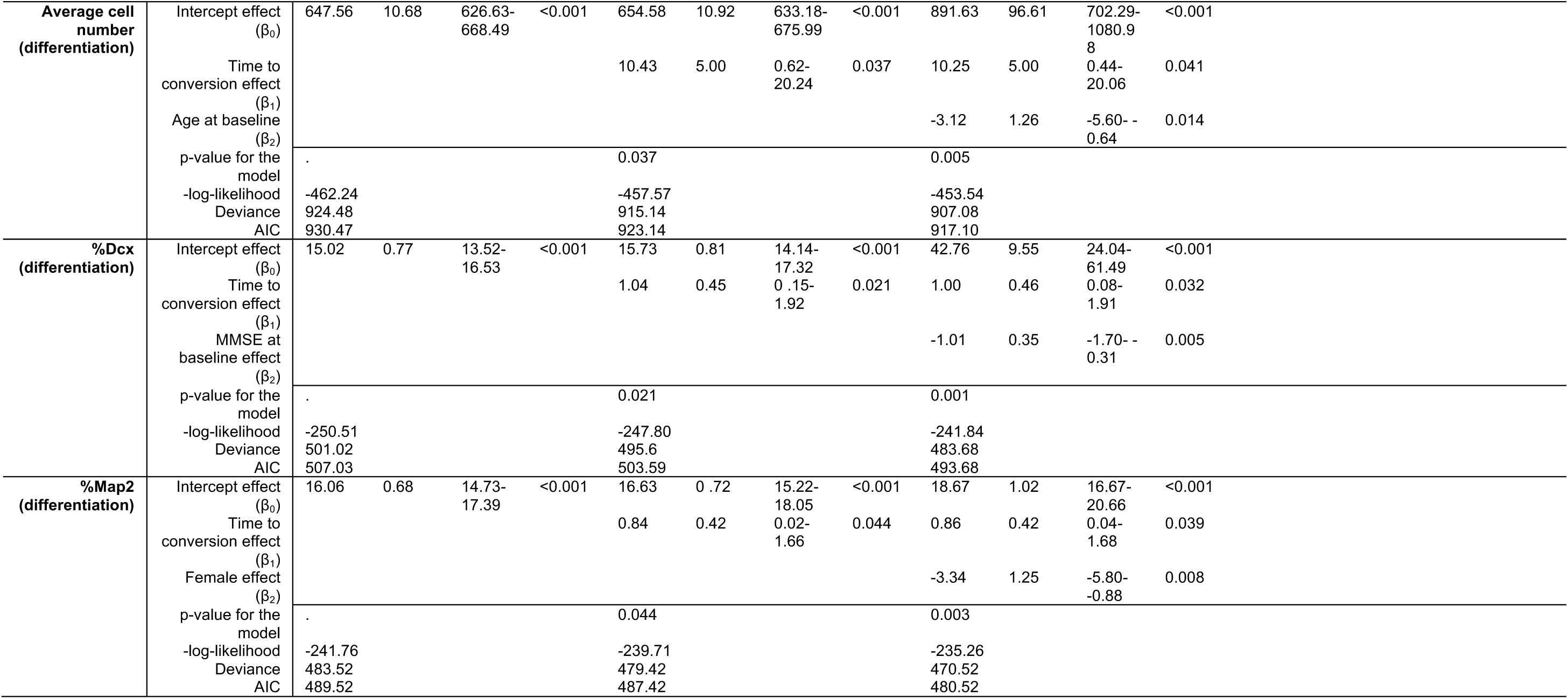
Factors affecting changes in the human hippocampal neurogenesis (dependent variable) when the HPC0A07/03C cells are exposed to 1% serum from the MCI converters. Models with the lowest Akaike Information Criterion (AIC) and deviance were selected as the best fit. Coefficient estimates (β), standard errors SE(β), 95% confidence intervals (CI) around the regression coefficient and significance levels for all predictors in the analysis are provided.

In addition, the mixed effects model with a random slope predicts that the level of apoptotic cell death (%CC3+ cells, p<0.0001, Fig.3, Table 1) increases with decreasing time to conversion to AD. For individuals with higher education that increase was smaller as compared to those with less than 10.5 years of education, suggesting that environmental factors related to education level, such as diet, might significantly impact on cell death during disease progression.

The percentage of variance that accounted for the average cell count was 66.9%, 88.92% for Ki67 and 88.3% for CC3, indicating excellent fit of the models.

### Increased hippocampal neurogenesis characterizes progression from MCI to AD

Human hippocampal progenitor cells were also exposed to the serum samples during differentiation to investigate neurogenesis (Fig. 4). Mixed effects models with random intercept (for cell count and %Dcx) or random slope (for %Map2) predict that there is an increased cell number (p=0.037, Table 1) and increased level of neuroblast (%Dcx, p=0.021, Table 1) and more mature neurons (%Map2, p=0.044, Table 1) with disease progression. We did not detect significant alterations in the number of apoptotic human hippocampal progenitor cells during differentiation.

**Fig. 4.**
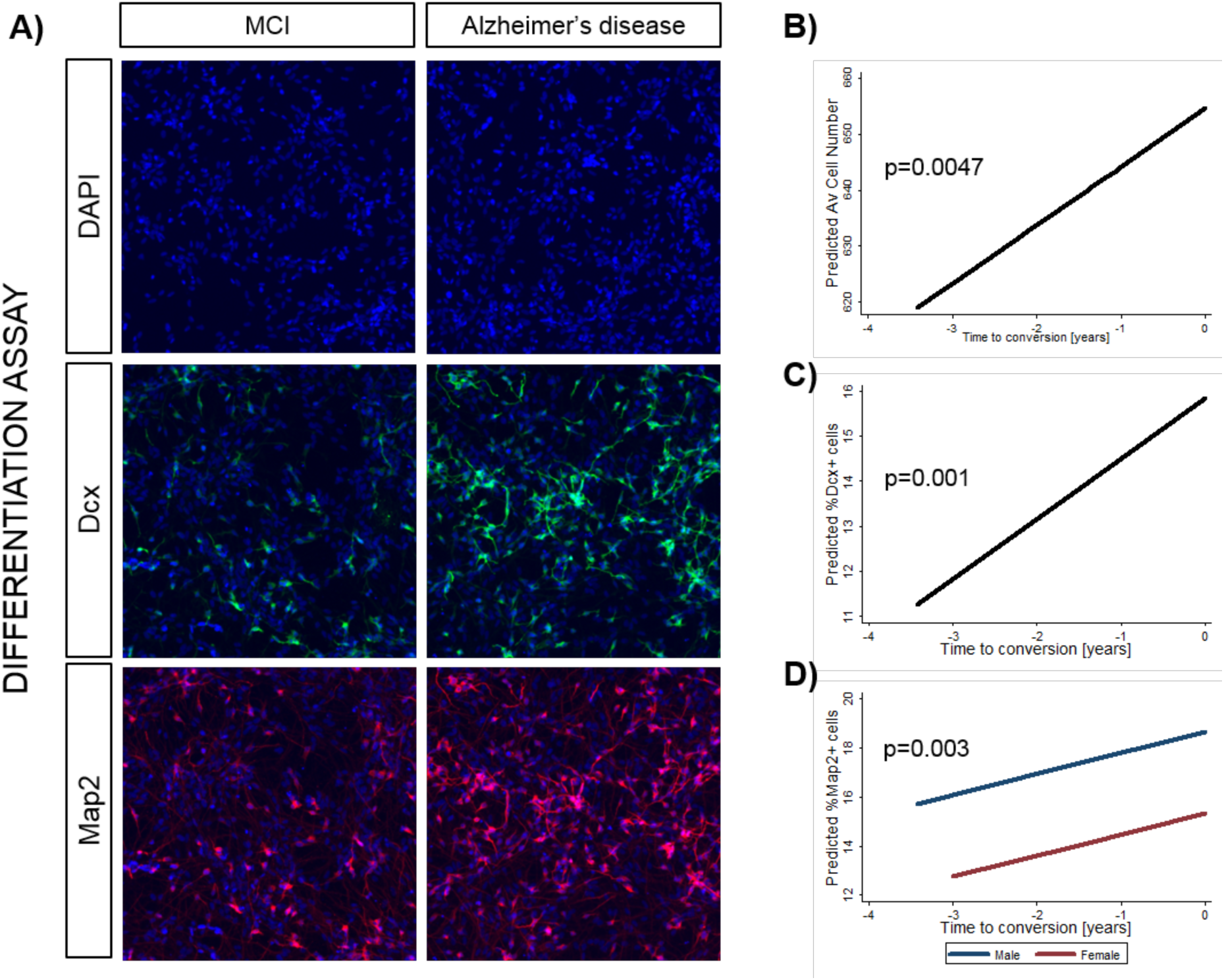
Exposure to 1% serum from the converters to Alzheimer’s disease leads to increased human hippocampal neurogenesis in vitro. **(A)** Representative images showing changes in human hippocampal progenitor cell number (DAPI) and neurogenesis (neuroblast marker doublecortin (Dcx) and neuronal marker microtubule-associated protein 2 (Map2)) when the HPC0A07/03C cells are cultured in the presence of 1% serum from an individual that converted to dementia due to AD (right panel, labelled Alzheimer’s disease) or serum from the same donor, collected 1 year before conversion (left panel, labelled MCI). **(B-D)** Trajectories of human hippocampal neurogenesis changes were fitted using mixed-effects models. Predicted mean scores are presented on the graphs, with the x axis representing time to conversion, with conversion assigned year=0 value and time before conversion assigned negative values for respective years. Longitudinal serum samples from MCI converters lead to **(B)** increased average cell count, **(C)** increased number of immature (%Dcx-positive cells) and **(D)** mature (%Map2-positive cells) neurons. All fitted regression lines were calculated from models presented in Table 1.

Time to conversion was the only significant predictor of the average cell number during the differentiation stage of the assay (see Table 1). Mini–Mental State Examination (MMSE) score at baseline and time to conversion predicted the number of immature neurons, i.e. the higher MMSE score at baseline, the lower number of Dcx-positive neurons. Gender of the serum donors was significantly influencing the predicted level of mature neurons, with females having on average 3.34% fewer Map2-positive cells as compared to males, when controlling for the time to conversion.

The percentage of variance that accounted for average cell count was 72.47%, 58.31% for Dcx, 55.83 % for Map2, indicating a moderate fit of the models.

Variables such as *APOE4* status and comorbidities (as listed in Table 2) were not significant when included in the mixed-effects models with respect to proliferation and differentiation stages of the assay. No significant interaction effects were determined.

**Table 2.** Baseline characteristics and epidemiological history of the study participants. Comorbidities represent either history of disease or being presently affected. Drug intake means either history of medications or current intake. All data is presented as number (%) or mean (±SD).

| **Baseline characteristics** | **MCI converters** | **MCI non-converters** |
| --- | --- | --- |
| Gender (%; female; F/M) | 60.52% | 38.89% |
| APOE4+ status (%) | 54.29% | 47.06% |
| Age at baseline (years±SD) | 76.02±7.81 | 78.05±5.08 |
| MMSE at baseline (±SD) | 26.78±1.97 | 27.94±1.43 |
| Education (years±SD) | 8.95±4.53 | 12.83±3.09 |
| Years from baseline until conversion/last assessment (±SD; range) | 2.35±0.79; 2-5.08 | 4.86±1.35; 2.92-7 |
| **Comorbidities (Objective and Self-reported)** | **MCI converters** | **MCI non-converters** |
| Hypertension | 12 (31.58%) | 10 (55.55%) |
| Heart attack ever | 1 (2.63%) | 4 (22.22%) |
| Angina | 4 (10.53%) | 3 (16.67%) |
| Depression | 20 (52.63%) | 9 (50%) |
| Arthritis | 1 (2.63%) | 3 (16.67%) |
| Asthma | 2 (5.26%) | 1 (5.55%) |
| Cancer | 5 (13.16%) | 6 (33.33%) |
| Glaucoma | 2 (5.26%) | 1 (5.55%) |
| Hypothyroidism | 2 (5.26%) | 3 (16.67%) |
| Infections, allergies | 5 (13.16%) | 2 (11.11%) |
| Stroke/TIA | 4 (10.53%) | 2 (11.11%) |
| Diabetes | 7 (18.42%) | 2 (11.11%) |
| **Drug intake** | **MCI converters** | **MCI non-converters** |
| Antidepressants | 13 (34.21%) | 3 (16.67%) |
| NSAIDs | 1 (2.63%) | 0 (0%) |
| Analgesics | 7 (18.42%) | 5 (27.78%) |
| AD drugs | 11 (28.95%) | 0 (0%) |
| Metformin | 4 (10.53%) | 1 (5.55%) |
| Sleeping pills | 3 (7.89%) | 2 (11.11%) |
| Statins | 15 (39.47%) | 8 (44.44%) |
| **Lifestyle factors** | **MCI converters** | **MCI non-converters** |
| Excessive alcohol intake | 1 (2.63%) | 3 (16.67%) |
| Smoking (ever) | 4 (10.53%) | 7 (38.89%) |
| Solitary living | 16 (42.10%) | 9 (50%) |
| Supplements* intake | 7 (18.42%) | 5 (27.78%) |

\* Vitamins (incl. folic acid) and omega-3 fatty acids

### Hippocampal cellular readout can distinguish MCI converters from MCI non-converters

Next, we asked whether serum from MCI converters compared to serum from MCI non-converters impacted differentially on proliferation and differentiation of human hippocampal progenitor cells. Using fitted models we demonstrated that during the proliferation stage of the assay the number of hippocampal progenitor cells exposed to serum increases with consecutive follow-up visits. Nonetheless, the number of cells treated with serum from MCI converters was higher as compared to the effect of serum from MCI non-converters (p<0.0001, Fig. 5, Table 3). On the other hand, we observed that the number of Ki67-positive cells decreases with time, however significantly less so when they were subjected to serum from MCI converters (p=0.0001, Fig. 5).

**Fig. 5.**
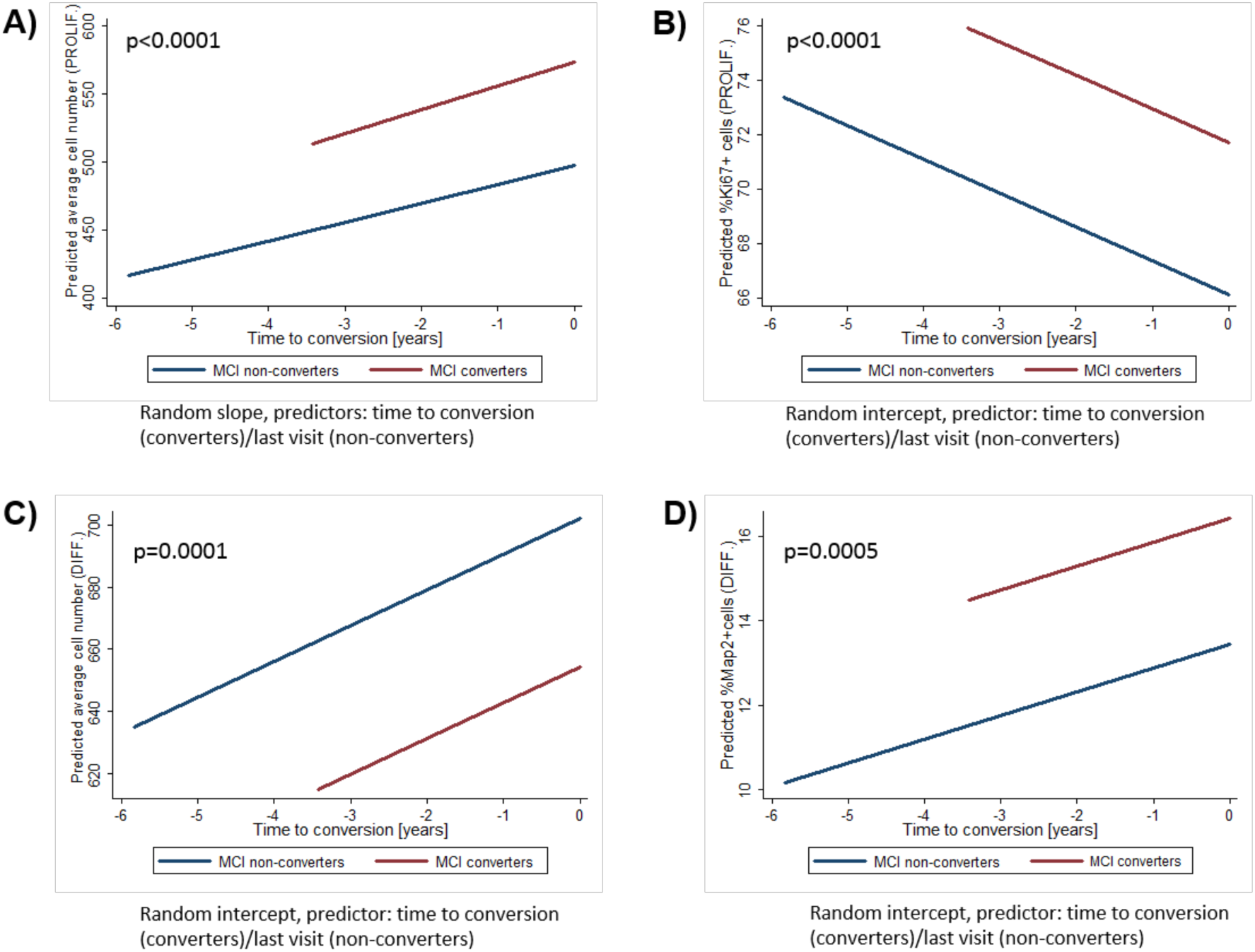
Significant differences in the effects of longitudinal serum samples from MCI converters and MCI non-converters on human hippocampal cells proliferation (A, B) and differentiation (C, D). Trajectories of human hippocampal neurogenesis changes with follow-up visits were fitted using mixed-effects models for repeated measures. Predicted mean scores are presented on the graphs, with the x-axis representing time to conversion (with conversion assigned year=0 value and time before conversion assigned negative values for respective years for the MCI converters) or time to last visit from baseline (for the non-converters). Longitudinal serum samples lead to higher average cell count during both proliferation **(A)** and differentiation **(C)**, as well as to smaller decrease in proliferation **(B)** and increased neurogenesis **(D)**. All fitted regression lines were calculated from models presented in Table 3.

**Table 3.** Significant factors affecting human hippocampal neurogenesis (dependent variable) when the HPC0A07/03C cells are exposed to 1% serum from either MCI converters or MCI non-converters. Models with the lowest Akaike Information Criterion (AIC) and deviance were selected as the best fit. Coefficient estimates (β), standard errors SE(β), 95% confidence intervals (CI) around the regression coefficient and significance levels for all predictors in the analysis are provided.

| Model outcome | Model measure | Random intercept (model 0) |  |  |  | Random intercept (model 1) |  |  |  | Random slope (model 2) |  |  |  |
| --- | --- | --- | --- | --- | --- | --- | --- | --- | --- | --- | --- | --- | --- |
| | | $\beta$ | SE ($\beta$) | 95% CI | p | $\beta$ | SE ($\beta$) | 95% CI | p | $\beta$ | SE ($\beta$) | 95% CI | p |
| Average cell number (proliferation) | Intercept effect ($\beta_0$) | 530.78 | 10.93 | 509.36-552.20 | <0.001 | 498.54 | 16.67 | 465.86-531.22 | <0.001 | 499.05 | 15.95 | 467.80-530.31 | <0.001 |
| | Conversion effect ($\beta_1$) | - | - | - | - | 74.79 | 19.83 | 35.92-113.65 | <0.001 | 74.90 | 19.16 | 37.35-112.44 | <0.001 |
| | Time to conversion/last visit effect ($\beta_2$) | - | - | - | - | 17.08 | 2.71 | 11.75-22.40 | <0.001 | 18.24 | 3.91 | 10.59-25.90 | <0.001 |
|  | p-value for the model | . |  |  |  | <0.0001 |  |  |  | <0.0001 |  |  |  |
|  | -log-likelihood | -900.10 |  |  |  | -867.74 |  |  |  | -864.25 |  |  |  |
|  | Deviance | 1800.2 |  |  |  | 1735.48 |  |  |  | 1728.5 |  |  |  |
|  | AIC | 1806.21 |  |  |  | 1745.49 |  |  |  | 1740.49 |  |  |  |
| Average cell number (differentiation) | Intercept effect ($\beta_0$) | 658.60 | 9.13 | 640.70-676.50 | <0.001 | 701.94 | 16.06 | 670.46-733.41 | <0.001 | | | | |
| | Conversion effect ($\beta_1$) | | | | | -47.64 | 18.98 | -84.83- -10.44 | 0.012 | | | | |
| | Time to conversion/last visit effect ($\beta_2$) | | | | | 11.48 | 2.95 | 5.69-17.27 | <0.001 | | | | |
|  | p-value for the model |  |  |  |  | 0.0001 |  |  |  |  |  |  |  |
|  | -log-likelihood | -889.37 |  |  |  | -874.64 |  |  |  |  |  |  |  |
|  | Deviance | 1778.74 |  |  |  | 1749.28 |  |  |  |  |  |  |  |
|  | AIC | 1784.73 |  |  |  | 1759.28 |  |  |  |  |  |  |  |
| %Map2 (differentiation) | Intercept effect ($\beta_0$) | 14.82 | 0.60 | 13.65-15.99- | <0.001 | 13.45 | 1.02 | 11.46-15.44 | <0.001 | | | | |
| | Conversion effect ($\beta_1$) | | | | | 2.96 | 1.19 | 0.64-5.29 | 0.012 | | | | |
| | Time to conversion/last visit effect ($\beta_2$) | | | | | 0.56 | 0.23 | 0.12-1.01 | 0.014 | | | | |
|  | p-value for the model | . |  |  |  | 0.0005 |  |  |  |  |  |  |  |
|  | -log-likelihood | -457.37 |  |  |  | -449.54 |  |  |  |  |  |  |  |
|  | Deviance | 914.74 |  |  |  | 899.08 |  |  |  |  |  |  |  |
|  | AIC | 920.73 |  |  |  | 909.08 |  |  |  |  |  |  |  |

In addition, serum from MCI converters leads to higher apoptotic cell death during both proliferation (p=0.0001, R^2^= 0.099, Adj R^2^= 0.093) and differentiation (p<0.0001, R^2^= 0.263, Adj R^2^= 0.258). This effect, however, is time-independent.

Fitted models indicated that during the differentiation stage of the assay there was an overall decrease in the number of cells treated with serum, and human hippocampal progenitor cells treated with serum from MCI converters were characterized by significantly higher drop in number compared to cells treated with serum from MCI non-converters (p=0.0001, Fig. 5, Table 3). Moreover, the number of mature neurons (Map2-positive cells) significantly increased when cells were treated with serum from MCI converters (p=0.0005, Fig. 5).

### Prediction of conversion to AD using baseline data

Our next research question was whether the baseline cellular readouts together with baseline patients’ data could predict conversion to AD. Using stepwise logistic regression, the best predictors of conversion from MCI to AD were determined, i.e. education (years) and cellular readouts from both proliferation and differentiation stages of the assay -average cell count during proliferation, %Ki67-positive cells and %CC3-positive cells during differentiation (Table 4).

**Table 4.** Predictors of conversion to Alzheimer’s disease from stepwise logistic regression analysis. Predictors: Education (measured in years), average cell count during proliferation, proliferation marker (Ki67), and apoptosis during differentiation (cleaved caspase 3, CC3). Model parameters: R2 =0.6672, p>0.0001.

| Predictor | Odds Ratio | P>z |
| --- | --- | --- |
| Education (years) | 0.72 | 0.040 |
| Average cell count (proliferation) | 1.03 | 0.001 |
| %Ki67+ cells (proliferation) | 1.35 | 0.038 |
| %CC3+ cells (differentiation) | 3.49 | 0.016 |
| Intercept | 1.68e-15 | 0.012 |

The fit of the model was confirmed using Hosmer-Lemeshov goodness of fit (p=0.324) and Stata linktest, demonstrating no specification errors (_hat=0.001, _hatsq=0.110). To assess the accuracy of the selected predictors in assigning individuals into MCI converters or non-converters groups, receiver operating characteristic (ROC) analysis was performed. ROC curve (Fig. 6) demonstrated excellent discriminatory power (area under the curve, AUC=96.75 %) of the model to distinguish between future converters and non-converters. The model correctly classified 92.11% converters and 94.12% MCI non-converters were correctly discriminated.

**Fig. 4.**
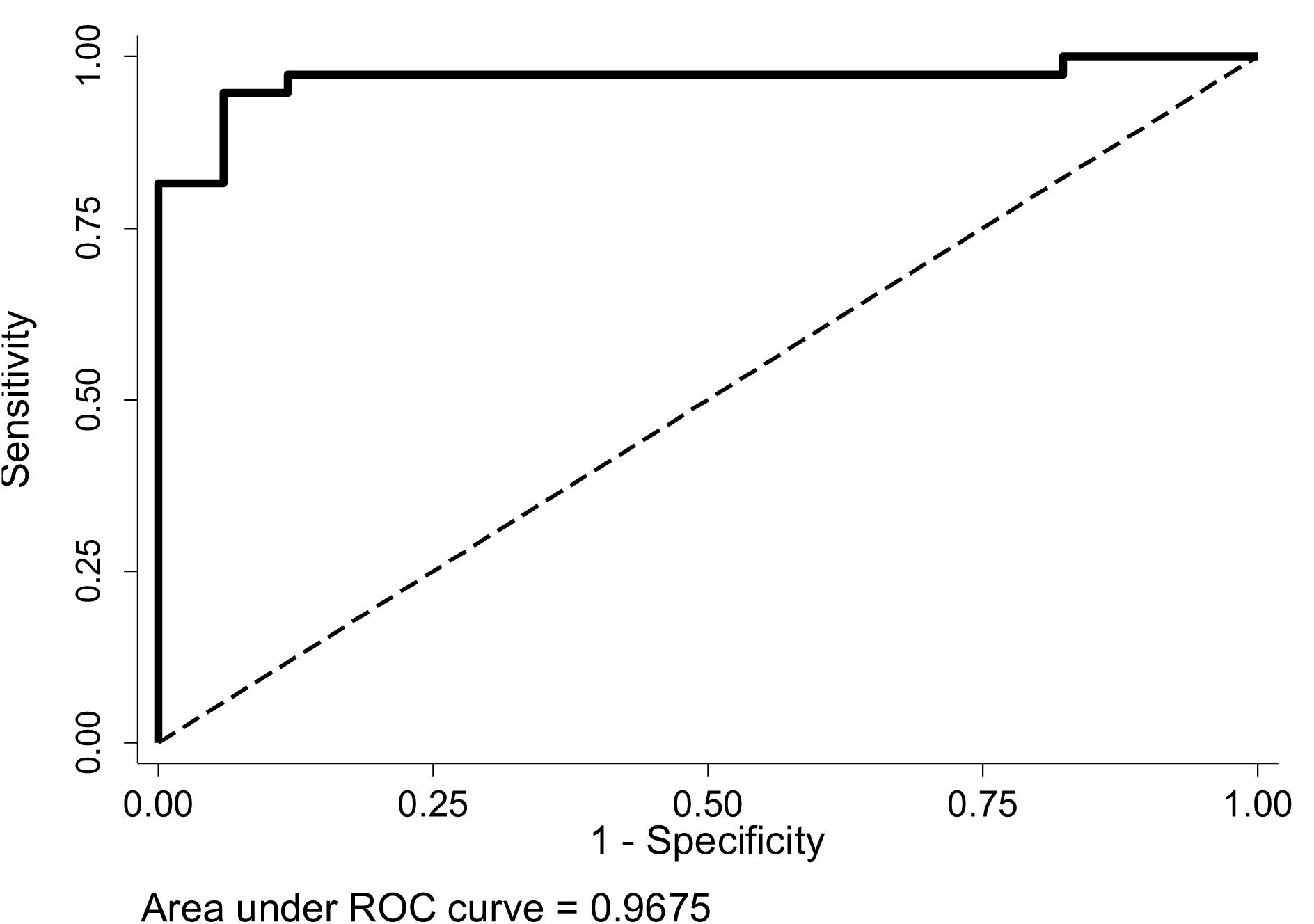
Receiver-operator characteristic-curve for predicting conversion to Alzheimer’s disease. The prediction model (AUC=0.9675) indicates an excellent discriminative performance of the model. Sensitivity = 92.11%, Specificity = 94.12%, Positive predictive value = 97.22%, Negative predictive value = 84.21%.

In supplementary Table S1, AUC for each predictor alone is presented, showing that the model built of all four predictors is characterized by better parameters.

### Machine learning cross-validation of the prediction model of conversion to AD

To validate the ROC curve described above, internal validation using machine learning was performed. Support Vector Machine Classifiers using the Radial-Based Kernel were trained to predict conversion status using data from the 38 MCI converters and 18 MCI non-converters. Four predictors were used, including the neurogenesis factors (average cell count during proliferation, %Ki67-positive cells and %CC3-positive cells during differentiation) and education. Performance of the classifier was assessed using 1000 repeats of 5-fold cross-validation. The ‘e1071’ package in R was used to train and test the classifiers. We found that a classifier using these three neurogenesis factors as predictors achieved an area under the curve of 0.93, with sensitivity 90.3% and specificity 79.0% (Fig. S1).

### Molecular pathways involved in hippocampal neurogenesis and AD differentially expressed in baseline serum of MCI converters and non-converters

A proteomic analysis using the SOMAScan assay followed by Ingenuity Pathway analysis was applied to gain insights into molecular pathways and networks by which proteins in serum might regulate the hippocampal stem cell fate and conversion to AD. The serum levels of 207 proteins (Table S2) were found to be significantly differentially expressed between MCI converters and non-converters. A classifier trained on 207 proteins was able to distinguish between serum samples from MCI converters and non-converters with excellent diagnostic performance (AUC=0.943), with sensitivity of 91.65% and specificity of 81.68% (Fig. S2).

Among the differentially expressed proteins between MCI converters and non-converters, there were proteins involved either in the neurogenic process (e.g. GDF11) or in AD (e.g. CA2, DKK1, MAPT), or in both (e.g. CREBBP, SFRP1, DKK1, APOE, IL1RAP).

We applied Ingenuity Pathway Analysis (IPA) to analyse the differentially expressed proteins between MCI converters and non-converters. The identified canonical pathways included p38 MAPK Signalling (p=3.28E-02, ratio 7/55), PPAR Signalling (p=3.48E-02, ratio 6/44), Wnt/β-catenin Signalling (p=5.48E-02, ratio 6/49), Granulocyte Adhesion and Diapedesis (p-value=2.82E-02, ratio 10/90), Agranulocyte Adhesion and Diapedesis (p=4.18E-02, ratio 9/83). The two top generated networks had a score of 36 (Cell-To-Cell Signalling and Interaction, Cellular Movement, Immune Cell Trafficking, and Cellular Movement, Cell Death and Survival, Embryonic Development, presented in Fig. 7). Both top networks were characterized by molecules central to HN and AD as their major hubs (NFkβ and Jnk in Cell-To-Cell Signalling and Interaction, Cellular Movement, Immune Cell Trafficking, and ERK1/2, ERK and p38 MAPK in Cellular Movement, Cell Death and Survival, Embryonic Development).

**Fig. 5.**
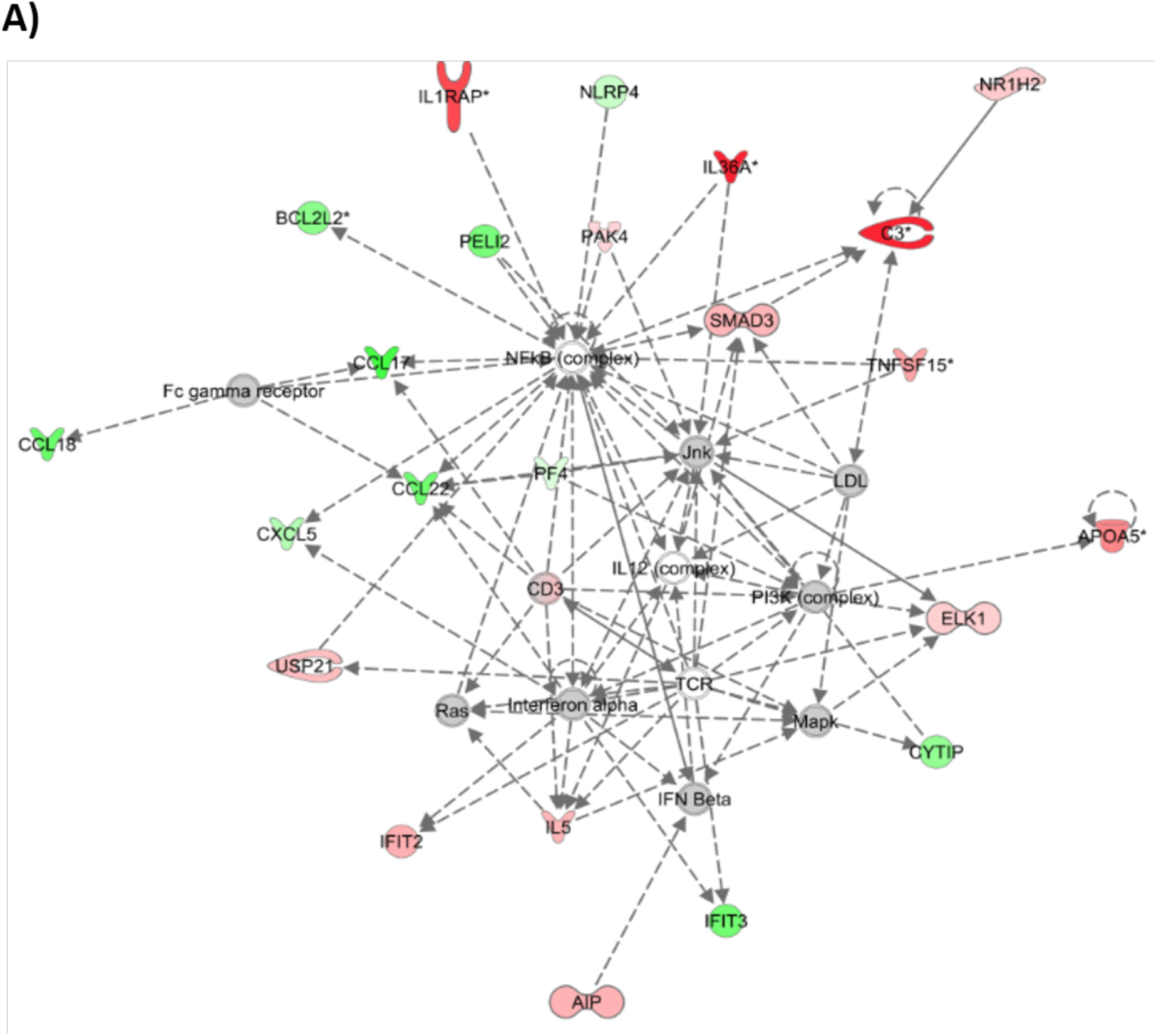
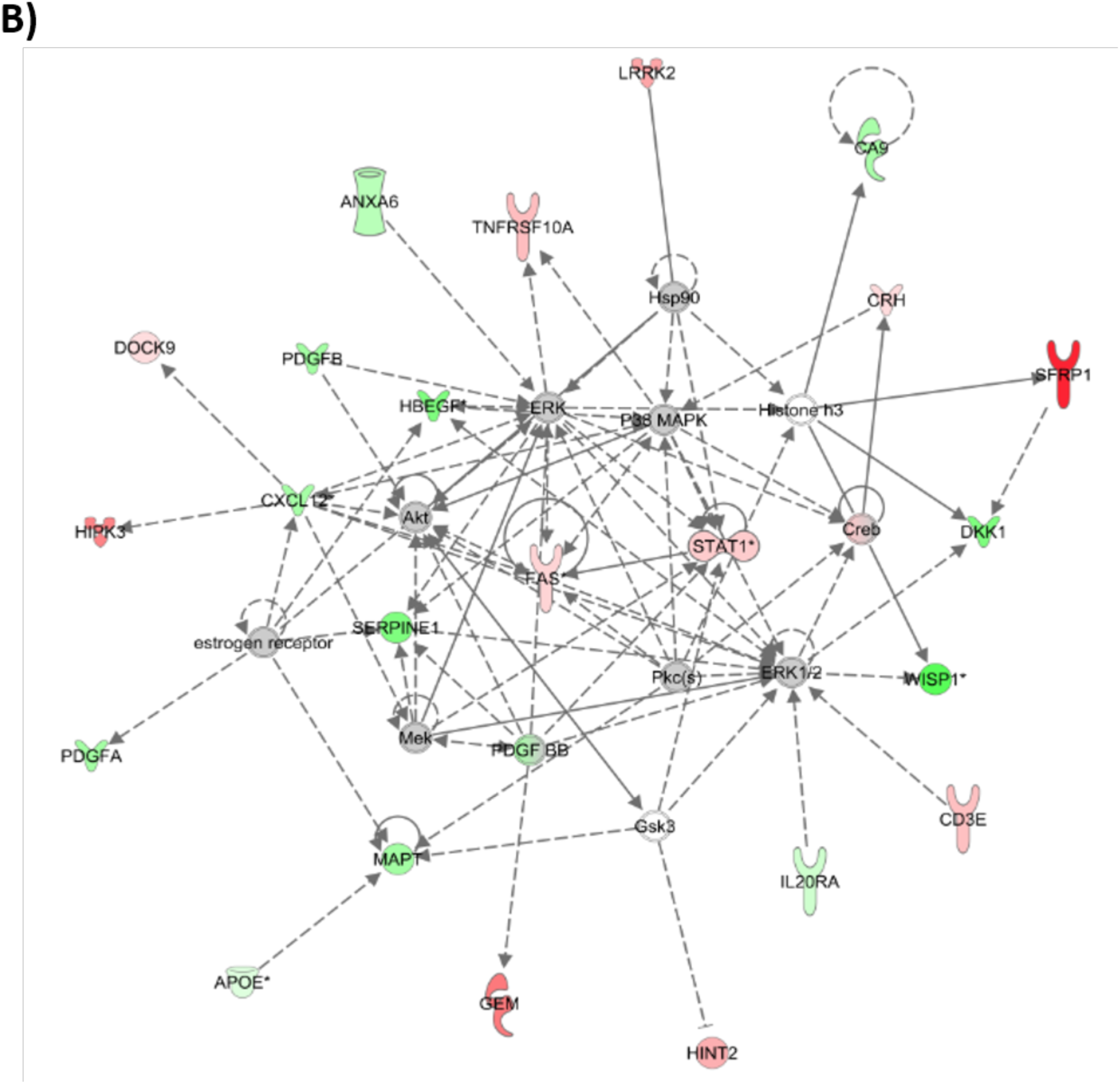
The two top scoring networks generated by IPA, Cell-to-Cell Signalling and Interaction, Cellular Movement, Immune Cell Trafficking (A); Cellular Movement, Cell Death and Survival, Embryonic Development (B). Networks were generated 207 differentially expressed proteins by MCI converters, out of which 199 were identified by IPA. **A)** Cell-to-Cell Signalling and Interaction, Cellular Movement, Immune Cell Trafficking network (score 36) comprises 23 focus molecules analysed in the SomaScan panel; **B)** Cellular Movement, Cell Death and Survival, Embryonic Development (score 36) comprises 23 focus molecules analysed in the SomaScan panel. Proteins are represented by nodes, with up-regulated proteins displayed in red, and the down-regulated proteins are in green, and additional interacting molecules not included in the SomaScan are marked in white. Each network is displayed as a series of nodes (proteins) and edges (i.e. lines, corresponding to biological relationships between nodes). The solid lines indicate direct interactions, whereas the dotted lines represent indirect interactions between the proteins represented in the networks.

## Discussion

This study has introduced a new “in vitro parabiosis” assay employing a human hippocampal progenitor cell line exposed to longitudinal human serum samples. This assay enables studying the effects of the changing systemic milieu on neural stem cells overtime. Using this unique model, we provide potential insights into temporal changes of the hippocampal neurogenic process during AD progression, with baseline data predicting with excellent accuracy conversion from MCI to AD up to 3.5 years before clinical diagnosis.

Previous in vivo rodent parabiosis experiments demonstrated significant role of the systemic environment in determining the fate of hippocampal progenitor cells (*10, 13*). More recently, Middeldorp et al. reported that plasma from young mice restored some aspects of hippocampal plasticity and associated learning in AD mouse models (*28*). Here, we show for the first time the ability of the MCI and AD human systemic environment to differentially regulate human neural stem cell fate along disease progression and conversion.

We report an increase in HN induced by serum obtained at around conversion time from MCI to AD. Human autopsy studies demonstrated dysregulation of HN in AD, however it has been debated whether HN is increased (*29*), decreased (*30*) or unchanged in AD (*31*). AD animal models brought similarly contradictory results (for review see (*7*)). Importantly, the majority of the autopsy and rodent reports describe HN at the late stage of AD, thus it is impossible to extrapolate these results to the early stage of the disease. Although our in vitro measures are only potential proxy of in vivo HN, our data is in line with a recent rodent study investigating HN in prodromal AD showing that proliferation of hippocampal DCX expressing neuroblasts was significantly and specifically elevated during the pre-plaque stage in the APP-PS1 AD model (*8*).

The functional role of HN is not simply to replace dying neurons, but instead, to add immature neurons to the existing circuits in the hippocampus and reshape these in response to new stimuli (*32*). The immature neurons do not only have distinct properties from those of mature neurons, but are also more plastic and excitable as compared to the mature granule neurons. HN plays a critical role in reducing interference between similar memories (pattern separation, temporal separation of memories (*33*)), conjunctive encoding (integrating spatial and non-spatial information into memory), consolidation of contextual memories (*34*), emotional regulation and in response/adaptation to stress (*4, 33-36*). On the other hand, decreased HN is associated with cognitive impairment (*37, 38*). It remains uncertain if and how changes in HN are relevant for human cognition, and we do not understand yet if increased HN plays a compensatory role, provides cognitive resilience or might contribute to the ongoing pathology. Despite the fact that increased neurogenesis after an injury such as stroke or traumatic brain injury improves cognition, it appears that in general it does not always result in an improved function. For instance, aberrant neurogenesis has been implicated in epilepsy pathogenesis (*39-42*). Neuronal hyperactivation leads to long-term impairment of HN, with an increase in neurogenesis followed by inappropriate migration, differentiation and integration of many of new neurons (*43, 44*).

Interestingly, increased hippocampal activity has been reported in MCI individuals and was not linked with significant hippocampal volume changes (*45-52*). This hyperactivation occurs in the DG and CA3 subfields of the hippocampus (*50*). Bakker et al. demonstrated that reducing hippocampal hyperactivation with an antiepileptic drug, levetiracetam, has a therapeutic potential as it improves pattern separation in the individuals with MCI (*52*). In the future, it would be interesting to investigate patterns of hippocampal activity alongside the presented in vitro assay, where changes in HN might correspond to hyperactivation in the individuals with MCI who will eventually convert to AD.

Interestingly, increased HN might interfere with retrieval of old memories (*34, 35, 53*) and ablation of neurogenesis improves hippocampal-dependant working memory by reducing interference (*54, 55*). For instance, it was reported that too much neurogenesis might be detrimental for the hippocampus-dependent working memory, involving both the hippocampus and the prefrontal cortex (*54*).

Further, the CaM/Tet-DTA mice which are characterized by a selective neuronal loss in the hippocampus, have increased neurogenesis and angiogenesis, correlating with behavioural recovery, suggestive of a compensatory response (*56*). Despite increased neurogenesis in this mouse model, cognitive deficits were ameliorated only in the young (6-month old) (*56*) but not old (14-month old) mice (*57*). That might suggest that during age-related neurodegeneration, as in AD, increased HN might not be sufficient for cognitive recovery. It is likely that a permissive systemic milieu is indispensable to complement increased HN to induce cognitive improvement. A balance between age-and neurodegeneration-related factors and decreasing with age pro-youthful circulatory molecules determines HN level, and our study demonstrates that this phenomenon extends beyond rodents.

The presented prediction model identifies MCI converters and non-converters using baseline serum samples, combining education and relevant cellular readouts. That is in line with Tang et al. recommendation that the best prediction models should contain factors across different variable categories (*58*). Education attainment, an important factor in the presented prediction model, coincides with brain development, thus it can be a significant deterministic factor in adulthood cognition. Further, our model is concordant with previous studies, showing that lower education is associated with higher risk of AD (*59*). We show that the odds of converting to AD decrease by factor 0.72 with each additional year of education (see Table 4).

We believe that education attainment is a proxy of lifestyle and this concept extends beyond just purely the formal years of education. It might impact choice of occupation, socioeconomic status and may affect exposure to AD risk factors. Several animal studies demonstrated that both diet and environmental enrichment are significant factors affecting HN (*4, 60*). Therefore, in the future, collecting detailed information about individual’s lifestyle activities such as social and cognitive engagement, physical activity and diet might turn out to provide even better prediction of HN than education.

We did not find a relationship between available lifestyle-related factors or drug intake and HN. Similarly, only in the analysis of AD converters over time, we observed that some demographic characteristics, such as years of education, age at baseline, gender, and neuropsychological assessment tests such as MMSE at baseline, were predictors for cellular readouts. However, when comparing MCI converters vs non-converters over time, none of the cognitive or demographic measures significantly influenced HN. In our analyses, we did not detect any role for the *APOE4* status. The role of *APOE4* in mediating hippocampal volume and rate of hippocampal atrophy is debatable (*61-63*), thus we were not alarmed that it was excluded from our models.

Our findings suggest that the baseline serum sample is sufficient to predict conversion to AD with excellent accuracy. We also demonstrate that a panel of 207 serum proteins is nearly as good in predicting conversion as the proposed model. The activation of one of the canonical pathways revealed by IPA, p38 MAPK, might partially explain the neurogenesis readouts observed in our study. Activation of this pathway might trigger inhibition of proliferation, induce apoptosis and stimulate differentiation (*64*). Although the 207-protein panel comprised proteins of well-described role in neurogenesis and AD, one must interpret its components with caution, given that serum will also contains important factors other than proteins.

We recognize that our study has limitations. First, AD diagnosis was clinical only and none of the study participants had a post-mortem AD diagnosis. Secondly, there was no neuroimaging data available for some of the follow-up visits and our study cohort did not encompass an additional AD cohort with hippocampal sparing. Thirdly, despite the MCI converter and non-converter groups being both composed of a combination of patients/samples drawn from two independent cohorts, and having cross-validated our model with machine learning, we will need to validate our model in a third independent cohort. Future studies should also explore the generalisability of our results to familial AD.

Furthermore, we recognize that we do not reconstitute the neurogenic niche in its entirety, and future experiments should see the expansion of this in vitro model to include other key players in AD such as microglia, or extend the duration of the assay to monitor synaptic formation and plasticity. Finally, the effects observed in vitro might not mirror those *in vivo*. Yet, the assay could be used as a platform to explore longitudinal changes in HN and predict conversion to AD. Given that there are no methods to quantify or image HN in living individuals, our assay presents a powerful tool.

All together, we demonstrate that factors in the human systemic environment (i.e. serum) modulate human hippocampal progenitor cell fate. This novel in vitro assay enables monitoring disease progression and predicting conversion to AD up to 3.5 years before clinical diagnosis. Education and baseline in vitro assay readouts (proliferation: average cell count, Ki67; differentiation: cell death) serve as predictors in the model (AUC=0.9675).

The proposed assay has potential to facilitate early diagnosis and stratification in clinical trials. Early diagnosis of AD enables earlier implementation of interventions aimed at delaying symptom progression and making decisions regarding lifestyle changes. Finally, we believe that by understanding further the mechanisms underlying hippocampal progenitor cell alterations one could identify novel targets to prevent or delay AD onset and to slow down disease progression.

## Materials and Methods

### Cell line

All experiments were performed using the Multipotent Human Hippocampal Progenitor/Stem Cell Line HPC0A07/03C (ReNeuron, UK), derived from the first trimester female foetal hippocampal tissue following medical termination and in accordance with UK and USA ethical and legal guidelines, and obtained from Advanced Bioscience Resources (Alameda CA, USA). HPC0A07/03C cells were conditionally immortalised by introducing c-myc-ERTAM transgene which enables them to proliferate indefinitely in presence of epidermal growth factor (EGF), basic fibroblast growth factor (bFGF) and 4-hydroxy-tamoxifen (4-OHT) (*65*). Removal of these factors induces spontaneous differentiation into neurons, astrocytes or oligodendrocytes (*15, 27, 66*).

### Serum samples

Serum samples were collected from 56 individuals initially diagnosed with MCI. Thirty-eight of 56 MCI patients later developed dementia due to AD (MCI converters, with a minimum of two up to five yearly follow-up visits with cognitive assessment and blood collection), whereas 18 did not progress either to AD or other disease, had transient memory problems but remained cognitively stable over a period of at least 3 years from diagnosis (MCI non-converters, with up to six yearly follow-up visits with cognitive assessment and blood collection).

The serum samples were sourced from two independent cohorts:

1. EU AddNeuroMed Consortium, a multi-centre European study (*67, 68*), with six participating medical centres: University of Kuopio (Finland), University of Perugia (Italy), Aristotle University of Thessaloniki (Greece), King’s College London (United Kingdom), Medical University of Lodz (Poland) and University of Toulouse (France). Consensus diagnosis was made according to published criteria (*21, 69, 70*). Clinical diagnosis was confirmed during consecutive follow-up visits.
2. King’s Health Partners-Dementia Case Register (KHP-DCR), a UK clinic and population based study (*67*). Diagnosis of probable AD was made according to (*69*) and (*70*) criteria. MCI was diagnosed according to (*21*) criteria. Clinical diagnosis was confirmed during consecutive follow-up visits.

Informed written consent was obtained from all serum donors or their carers according to the Declaration of Helsinki (1991) and protocols and procedures were approved by the relevant Institutional Review Board at each collection site.

Longitudinal serum samples from the study participants were used to obtain a cellular readout corresponding to each follow-up visit and to analyse within-and between-subjects variability in how serum collected at different time-points affects HN in vitro. A minimum of three serum samples, collected at annual assessment, were required for MCI non-converters, and two samples, one prior to conversion and one post-conversion, for MCI converters. Serum was collected at the time of cognitive assessments and the samples were stored at −80°C, and underwent one freeze/thaw cycle before performing experiments.

### Baseline and longitudinal characteristics of serum donors

Baseline characteristics of serum donors are presented in Table 2. All participants were age-(p=0.320) and gender-matched (p=0.129, see Table 4). MCI converters completed significantly fewer years of education compared with MCI non-converters (p=0.002). They also scored significantly lower in MMSE (p=0.031). There was no difference in the number of the *APOE4* allele carriers (*APOE4*+) between MCI converters and MCI non-converters (p=0.625).

### In vitro assay to investigate the impact of the systemic environment on human hippocampal neurogenesis

To study how the systemic environment (i.e. serum) influences human HN, we developed and optimized an in vitro assay, in which HPC0A07/03C cells (see (*15*) for medium composition) were treated with 1% serum. Control conditions consisted of medium + 1:100 PenStrep (Thermo Fisher Scientific). Serum was added to the cell culture during both proliferation (48 hours) and differentiation (7 days) (Fig.2). Proliferation medium comprised 4-hydroxytamoxifen (4-OHT), epidermal growth factor (EGF) and basic fibroblast growth factor (FGF). 24 hours after seeding, cell medium was replaced with medium containing 1% serum. To analyse proliferation markers, cells were fixed in 4% paraformaldehyde after the total of 72 hours of cell proliferation.

In order to investigate systemic effects on differentiation of HPC0A07/03C cells, after 48 hours of proliferation in the presence of 1% serum (as described above), cells were enabled to spontaneously differentiate by culturing them in medium devoid of 4-OHT, EGF and FGF. Experiments were terminated after 7 days of differentiation to avoid medium change and another serum supplementation. In addition, one week of differentiation is sufficient to detect changes in the number of immature and mature neurons (*15, 27, 66*). Medium supplemented with 1% serum was added only once during both proliferation and differentiation of the HPC0A07/03C cells. Serum samples were never pooled. For each experiment three biological replicates (cells of three different passage numbers) were used and for each biological replicate there were technical triplicates. All experiments were performed using the HPC0A07/03C cells of passage number ranging from 15 to 24.

### Immunohistochemistry and semi-automatic analysis

Rabbit polyclonal anti-Ki67 (Abcam) antibody was used to assess proliferation; rabbit monoclonal anti-CC3 to assess apoptosis; rabbit polyclonal anti-Dcx (Abcam) and mouse monoclonal anti-Map2 (Abcam) to assess neuronal differentiation. Secondary antibodies were conjugated with Alexa 488 or Alexa 555 (Invitrogen) and nuclei were counterstained with DAPI. Semi-automatic quantification of cell numbers and phenotypes was performed in 96-well plates, using the high-throughput instrument Cellinsight Personal Cell Imager CX5 (ThermoFisher Scientific).

### Somascan analysis

3620 unique proteins or 4006 different protein epitopes were quantified using the SOMAScan assay (SomaLogic Inc.,Boulder, CO, U.S.A.) from 150 ul of baseline serum samples. The SOMAScan assay is an aptamer-based technology that uses protein-capture SOMAmers (Slow Off-rate Modified Aptamer) to quantify proteins in a biofluid. SOMAmers are chemically modified oligonucleotides with specific affinity to their protein targets, developed by SELEX. The identities of all proteins quantified are listed in Supplementary Appendix 1 and the SOMAScan assay is described in detail at <u>www.somalogic.com.</u> The normalized and calibrated signal for each SOMAmer reflects the relative amount of each cognate protein present in the original sample, quantifications are reported in relative fluorescence units (RFU) and all data was first log10 transformed prior to analysis.

### Ingenuity pathway analysis

IPA (Ingenuity Pathway Analysis, IPA®, QIAGEN Redwood City) generated a list of canonical pathways and networks for proteins within detection limit of the SomaScan. Only proteins differentially expressed with p<0.05 were considered for analysis.

### Statistical analysis

#### Analyses of cellular readouts

All statistical analyses were performed using Prism 5.0 software (GraphPad Software), R or STATA 13. For univariate analyses two-tailed paired t-student test, one-and two-way ANOVA with post hoc comparisons using the Bonferroni correction were used. The chi-square test was carried out to test for differences in categorical outcomes such as gender and the *APOE4* allele.

Due to the longitudinal aspect of our dataset, we used linear mixed-effects regression models for repeated measures as they enable inclusion of varying number of assessment information available for each individual and do not require equal time intervals between the follow-up visits. Random intercept and random slope models were fitted with restricted maximum likelihood as the method of estimation. All analyses were adjusted for the effect of age, gender, education and carrying *APOE4* allele(s).

Each serum donor was assigned an ID in order to specify random effects in the models. The age of the individuals was centred at the cohort median (77 years) to aid interpretation of the models. Time to conversion was measured in years and centred at zero, which indicated conversion from MCI to AD. Time before conversion was assigned negative values and time after conversion positive values. *APOE4* allele carrier status was dichotomized – carriers of at least one *APOE4* allele were assigned 1, carriers of other APOE alleles – 0. Education was entered in the models either as years of education or it was dichotomized at the median (low ≤10, high >10 years of education). Classification to MCI converters or non-converters was dichotomous (MCI converters were assigned 1, non-converters −0).

Given that many individual characteristics or comorbidities might affect HN and/or AD risk, among the potential predictors considered in the models were: AD risk factors (gender, *APOE4* status, age centred around median and time to conversion (or time from baseline for MCI non-converters)), education level, solitary living and MMSE score (baseline MMSE, MMSE score change/year); comorbidities (diabetes, arthritis, hypertension, hypothyroidism, depression, cancer, stroke, angina, infections and allergies); drug intake (antidepressants, statins, nonsteroidal anti-inflammatory drugs); dietary supplements (vitamins, omega-3 fatty acids); life-style related factors (alcoholism, smoking). In addition, we also tested some biologically plausible interactions of different predictors. All mixed-effects regression models were assessed using Akaike Information Criterion (AIC), likelihood-ratio test and deviance.

#### Logistic regression prediction

Stepwise logistic regression analysis was carried out to assess the effect of selected predictors on probability of conversion to AD by 3.5-year follow up from baseline. It was preceded by analysis of multicollinearity. Receiver operating characteristic (ROC) curve was conducted to determine the classification accuracy of selected variables in predicting conversion to AD. Area under the curve was used to estimate the discrimination between MCI converters and non-converters.

#### Machine-learning classification

The ‘e1071’ package in R was used to train and test the machine learning classifiers based upon Support Vector Machine using the Radial Based Kernel. ROC curves were drawn using the “ROCR” package in R. ROC curves were drawn using the “ROCR” package in R. Performance of the classifier was assessed using 1000 repeats of 5-fold cross-validation.

## Supplementary Materials

**Fig. S1.** Receiver-operator characteristic-curve for the cross-validation model predicting conversion to Alzheimer’s disease.

**Fig. S2.** Receiver-operator characteristic-curve for predicting conversion to Alzheimer’s disease using a panel of 207 proteins.

**Table S1**. Comparison of AUC for logistic regression models for individual predictors.

**Table S2.** 207 proteins significantly differentially expressed between MCI converters and non-converters.

## Acknowledgements

We would like to thank the many research teams across Europe collecting samples for this and other AddNeuroMed studies’. We would also like to thank Rufina Leung for managing sample collection and distribution at the London site. AD acknowledge financial support from the National Institute for Health Research (NIHR) Biomedical Research and from the NIHR Collaboration for Leadership in Applied Health Research and Care South London at King’s College Hospital NHS Foundation Trust CLAHRC. The views expressed are those of the authors and not necessarily those of the NHS, the NIHR or the Department of Health

## Author contributions

AM and ST designed the experiments and wrote the paper. AM, TM, CDL carried out the in vitro cellular experiments, data collection and analyses. AM, BL, AD, AJN carried out the statistical analyses and the prediction models. BL and AJN performed the machine learning cross-validation. AM and BL carried out the proteomics analyses. CET and PJV collected patients’ samples and data. AM, JP, SL and ST designed the study. ST conceptualized the study. All authors read and revised the manuscript.

